# Ecological consequences of urbanization on a legume-rhizobia mutualism

**DOI:** 10.1101/2021.01.24.427992

**Authors:** David Murray-Stoker, Marc T. J. Johnson

**Affiliations:** Department of Ecology and Evolutionary Biology, University of Toronto, Toronto, Ontario M5S 3B2, Canada; Department of Biology, University of Toronto Mississauga, Mississauga, Ontario, Canada L5L 1C6; Centre for Urban Environments, University of Toronto Mississauga, Mississauga, Ontario, Canada L5L 1C6

**Author notes:** Corresponding author: David Murray-Stoker.

**Keywords:** mutualism, nitrogen, stable isotopes, urban ecology

## Abstract

Mutualisms are key determinants of community assembly and composition, but urbanization can alter the dynamics of these interactions and associated effects on ecosystem functions. Legume-rhizobia mutualisms are a model interaction to evaluate the ecological and ecosystem-level effects of urbanization, particularly urban-driven eutrophication and nitrogen (N) deposition. Here, we evaluated how urbanization affected the ecology of the mutualism between white clover (*Trifolium repens*) and its rhizobial symbiont (*Rhizobium leguminosarum* symbiovar *trifolii*) along an urbanization gradient. We found that the abundance of rhizobium nodules on white clover decreased with urbanization. White clover acquired N from mixed sources of N fixation and uptake from the soil for the majority of the urbanization gradient, but white clover primarily acquired N from the soil rather than N fixation by rhizobia at the urban and rural limits of the gradient. Importantly, we identified soil N as a critical nexus for urban-driven changes in the white clover-rhizobium mutualism. Taken together, our results demonstrate that urbanization alters the ecological consequences of a legume-rhizobium mutualism, with direct and indirect effects of the urban landscape on an ecologically-important mutualistic interaction.

## Introduction

Urbanization is a major driver of ecosystem change at local and global scales, consistently altering the ecological setting in terms of both biotic and abiotic factors (Grimm et al. 2008, Seto et al. 2010). Biotic changes frequently include fragmented habitats, homogenization of species composition, reduced abundance and diversity of native species, reduced vegetation cover, and increased abundance of non-native species (McKinney 2002, Grimm et al. 2008, Groffman et al. 2014, Aronson et al. 2016). Common changes to the abiotic environment include increased impervious surface cover, elevated temperatures, higher pollution levels (e.g., air, water, light, noise), and increased nutrient deposition (Grimm et al. 2008, Groffman et al. 2014, Stevens et al. 2018). However, there is limited research regarding the direct and indirect impacts of biotic and abiotic changes on species interactions in urban environments (Youngsteadt et al. 2015, Miles et al. 2019). Elucidating how urbanization affects the ecological consequences of species interactions is important for understanding the drivers of biodiversity and ecosystem change in urban environments.

Species interactions are key determinants of community composition (Wisz et al. 2013, Leibold and Chase 2017), but urbanization can disrupt these interactions (Raupp et al. 2010, Aronson et al. 2016, Miles et al. 2019) and their associated ecosystem functions (Ziter 2016). In urban landscapes, natural habitats are frequently fragmented and degraded, which can result in reduced species diversity and altered community composition (Williams et al. 2009, Aronson et al. 2016). Such ecological changes can alter antagonistic interactions (e.g., predator-prey, host-parasite; Rocha and Fellowes 2018, Parsons et al. 2019), competition (De León et al. 2019, Thomson and Page 2020), and mutualisms (Irwin et al. 2014, Rocha and Fellowes 2018, Rivkin et al. 2020). Mutualisms can be important for community and ecosystem stability, and disruptions to these interactions caused by urbanization may be particularly problematic for maintaining ecosystem functions. For example, pollinator abundance, diversity, and composition often change with urbanization (Harrison et al. 2018, Santangelo et al. 2020), which can shift the balance of the benefits conveyed between interacting plants and pollinators (Irwin et al. 2014, Rivkin et al. 2020, Santangelo et al. 2020). As another example, plant-microbe interactions are important for community assembly and nutrient cycling (van der Heijden et al. 2008), and urbanization can affect these interactions through altered soil chemistry mediated by pollution and nutrient deposition (Grimm et al. 2008, Stevens et al. 2018). As plant-microbe interactions are frequently nutrient-provisioning mutualisms, pollution and nutrient deposition that cause changes in the diversity and composition of microbial communities can have cascading effects on nutrient cycles in urban ecosystems (Galloway et al. 2003, Kaye et al. 2006). More broadly, if urbanization frequently alters mutualistic interactions, this may have cascading effects on communities and ecosystems.

Mutualisms between legumes and rhizobia are an ideal system for evaluating the ecological impacts of urbanization on species interactions. In these mutualisms, rhizobia fix atmospheric nitrogen in exchange for photosynthate and housing in nodules by their host plant (Hirsch 1992, Poole et al. 2018). Urbanization can disrupt these interactions, specifically through nitrogen (N) deposition and enrichment (Grimm et al. 2008, Zhang et al. 2012). Nitrogen deposition can inhibit the formation of nodules (Streeter and Wong 1988, Omrane and Chiurazzi 2009), precluding the development of the mutualism or reducing plant reliance on rhizobia for providing N (Vergeer et al. 2008, Weese et al. 2015,Regus et al. 2017). Additionally, N deposition can reduce N fixation rates by rhizobia (Cleland and Harpole 2010,Zheng et al. 2019). Although N deposition and short-term application of nutrient-rich fertilizers can benefit both legumes and rhizobia (Simonsen et al. 2015, Forrester and Ashman 2018), chronic and long-term exposure to increased N can reduce the ecological benefits of N fixation and cause the evolution of less beneficial rhizobia (Weese et al. 2015, Regus et al. 2017). Using the legume-rhizobia mutualism as a model system, it is possible to study how urbanization and nutrient deposition alter interactions in an ecologically-important mutualism.

In this study, we evaluated the hypothesis that urbanization alters the ecological and ecosystem-level consequences of a nutrient-provisioning mutualism. We used the mutualism between the legume white clover (*Trifolium repens*) and its rhizobial symbiont (*Rhizobium leguminosarum* symbiovar *trifolii*) as a tractable model interaction. We conducted our study along an urbanization gradient in Toronto, Canada (Fig. 1). Our study focused on three primary questions: (Q1) does rhizobia nodulation vary along an urbanization gradient? (Q2) How does the source of plant nitrogen (i.e., from soil or gaseous N_2_ fixed by rhizobia) change along the urbanization gradient? And (Q3) how do urban landscape features influence the interactions between soil N, plant N, and rhizobia nodulation? We predicted that: (1) urbanization would alter investment in rhizobia by *T. repens*, causing increased nodulation with decreased urbanization; (2) increased N availability in the soil due to urbanization would reduce N fixation by rhizobia and alter the source of N for *T. repens*; and (3) the changes in landscape features associated with urbanization cause direct and indirect effects on ecosystem structure and the white clover-rhizobium mutualism (Fig. 2).

**Figure 1.**
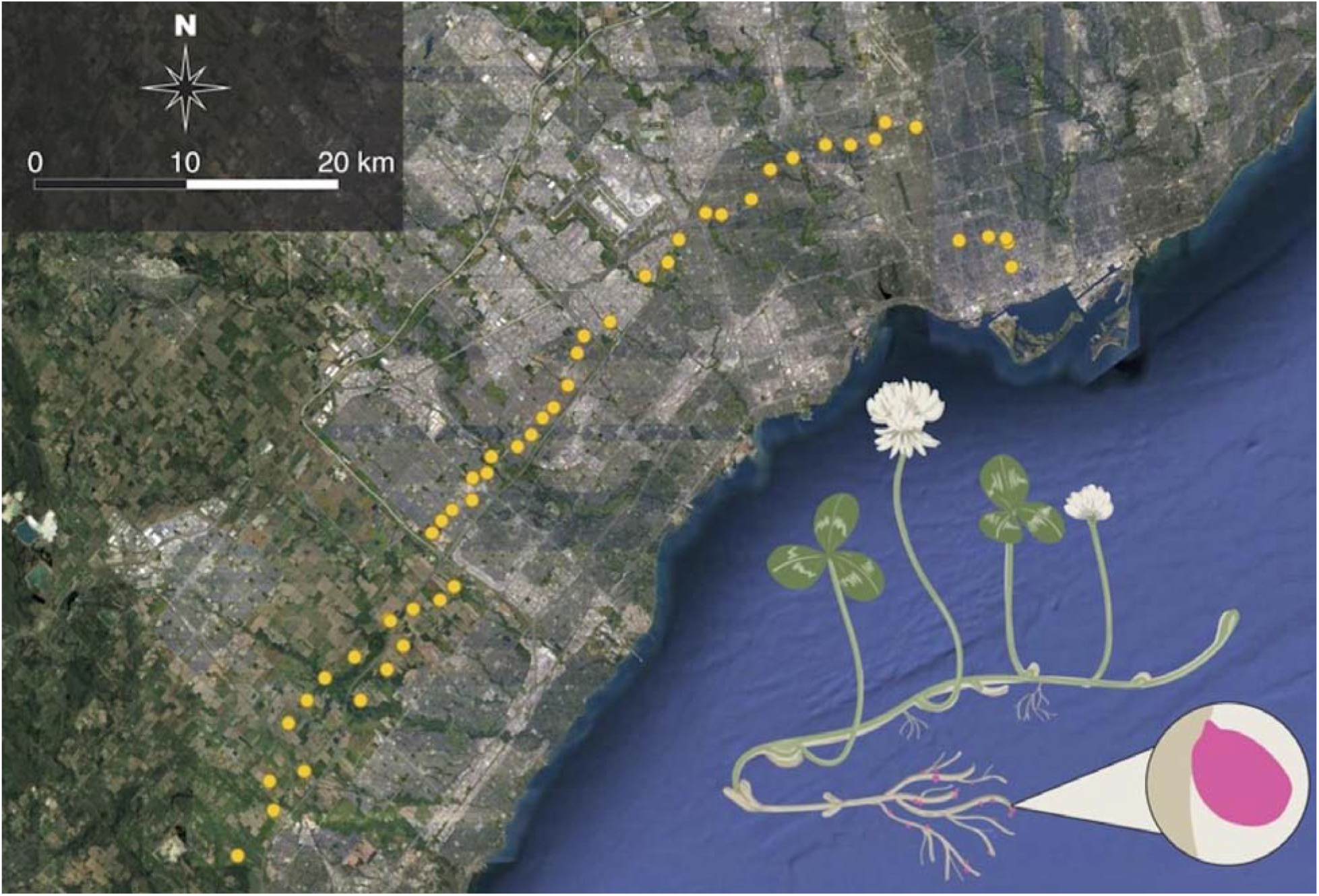
Map of the urbanization gradient in the Greater Toronto Area, ON, Canada displaying all 49 sampling sites for the study. The inset displays an illustration of a typical white clover individual sampled in the field, and the callout depicts a nodule attached to the root. Satellite imagery was taken in 2018 and retrieved from Google.

**Figure 2.**
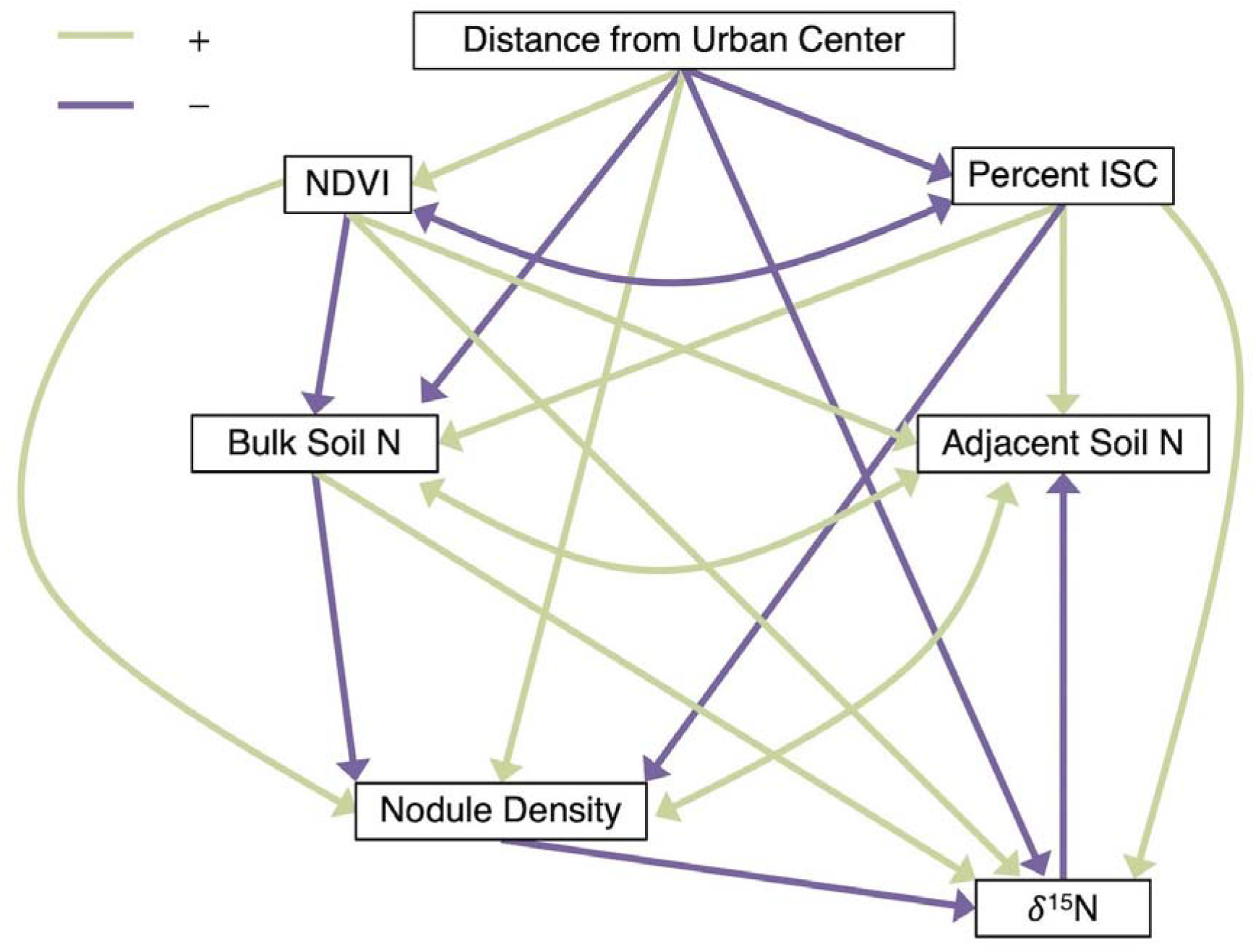
Hypothesized path model specifying causal and correlational pathways and the associated direction of each relationship. The structural set of linear equations for each causal or correlational pathway can be described by variables linking the response to the predictor (e.g., bulk soil N ∼ percent ISC + NDVI + distance). Distance from the urban center was fitted to account for extraneous sources of urbanization and environmental variation not explained by other predictors. We hypothesized that changes to local environments in the urban landscape would manifest in direct effects of urbanization (i.e., percent ISC and NDVI) on both soil N and the legume-rhizobia mutualism (i.e., nodule density and white clover δ^15^N), with indirect effects of urbanization mediated through soil N.

## Materials and Methods

### Land acknowledgement

We conducted our sampling at 49 sites along an urbanization gradient in the Greater Toronto Area of Ontario, Canada in August 2018 (Fig. 1). Our sampling was conducted on the traditional land of the Huron-Wendat, the Seneca, and most recently, the Mississaugas of the Credit First Nation.

### Study organisms

White clover (*Trifolium repens* L., Fabaceae) is a perennial, herbaceous legume native to Eurasia that is now globally distributed (Baker and Williams 1987). White clover typically reproduces clonally through stolons (Kemball and Marshall 1995) as well as through seed via obligately-outcrossed flowers (Barrett and Silander 1992). *Rhizobium leguminosarum* symbiovar *trifolii* is the primary rhizobial symbiont of *T. repens* (Martínez-Romero and Caballero-Mellado 1996, Andrews and Andrews 2017). As a facultative symbiont, *R. l*. bv. *trifolii* fixes atmospheric N_2_ into accessible NH_3_ for white clover in exchange for photosynthate and hosting within root nodules.

Nodulation is initiated when rhizobia attach to and penetrate the root. After infection of the root they continue to divide within the plant host cells to form nodules (Poole et al. 2018). At this stage, rhizobia differentiate into bacteroids and begin the N fixation process, whereby atmospheric N_2_ is converted into NH_3_ by bacteroids with the aid of nitrogenase enzymes (Hirsch 1992, Poole et al. 2018). Urbanization and specifically N enrichment may inhibit nodulation at multiple steps of the process. For example, NO_3_ and NH_3_ can inhibit rhizobia infection of roots, the formation of nodules, or the activity of nitrogenase (Streeter and Wong 1988, Omrane and Chiurazzi 2009). Another common mechanism for reduced nodulation with increased N availability is plant hosts switching N sources from N fixed by rhizobia to N taken up directly from the soil, resulting in lower nodulation and less investment in the symbiosis (Heath et al. 2010, Regus et al. 2017).

### Field sampling

Field collections of white clover, associated rhizobia, and soil samples were collected at all 49 sites along the urbanization gradient (Fig. 1). At each site, 9-10 clover individuals and associated roots and nodules were collected (mean ± SE = 9.94 + 0.13 individuals). Twenty soil cores were also taken at each sampling location, with 10 cores taken immediately adjacent to collected white clover and 10 taken at least 5 m from the nearest white clover plant (hereafter “bulk” cores), with the requirement that bulk cores did not contain other nitrogen-fixing legumes (e.g., *Medicago lupulina*). Each soil core was taken to a depth of 5 cm, and the top 2 cm plus organic material were removed because they could confound estimates of soil nutrients. Each sample type (bulk and adjacent) was combined for a composite sample of each soil core type per site, which were stored at −80°C until subsequent processing.

### Lab processing

#### Plant processing and nodule quantification

White clover samples were separated into leaf and root tissue. Leaf tissue was collected for stable isotope analyses by cutting 6-10 fully expanded, green, non-senescing leaves from each individual, which were placed into a 2 mL tube and stored in a freezer at -20°C until later processing. Root tissue was collected by cutting the first five roots below the plant base and directly attached to the stolon. Roots were measured to the nearest 1 mm until at least 10 cm of root length were measured or all collected roots were measured (root length: mean ± SE = 10.3 ± 0.13, range = 1.8-17.7 cm). Nodules were counted visually for each measured root, and counts were standardized by the total length of root measured to generate an estimate of nodule density per cm of root per plant.

#### Soil sample processing

Soil samples were prepared for N analyses by filtering, grinding, homogenizing, and drying samples. Samples were taken out of storage in the freezer at -80°C and thawed overnight in the lab (21.5°C). Each sample was then filtered over stacked sieves (mesh diameters: 4.75 mm, 2 mm, 1 mm, and 0.5 mm) to remove large rocks and gravel. Soil retained on the 0.5 mm sieve and catch pan was collected, and then samples were dried and stored at 60°C for 48 hrs. Samples were homogenized into a fine powder using mortar and pestle and stored in a drying oven at 60°C for approximately 4 weeks until later processing. Adjacent soil N provided an estimate of N available to white clover individuals and potentially affected by N-fixing rhizobia, and bulk soil N was a measure of background N availability at the site; adjacent and bulk soil were measured separately.

#### Leaf processing

Leaf samples were prepared for N analyses by freeze-drying, grinding, and drying the tissue. A composite sample for each population was prepared by taking one leaf from each of the white clover individuals per site and the composite samples were freeze-dried and then homogenized using a tissue grinder (FastPrep 96, MP Biomedicals, Irvine, CA, USA). Samples were dried and stored in a drying oven to achieve a constant mass at 60°C for 24-48 hrs until later processing.

#### Soil and plant N analyses

Soil and plant samples were weighed (soil ≈ 30 mg, plant ≈ 2 mg) on a micro-balance (XP2U Mettler Toledo, Mississauga, ON, Canada) and packed into aluminum capsules (Costech Analytical Technologies Inc., Valencia, CA, USA). Soil samples were analyzed for % N using a Carlo Erba NA 1500C/H/N Analyzer (Carlo Erba Strumentazione, Milan, Italy). Plant tissue was simultaneously analyzed for % N and N isotopes (i.e., ^15/14^N) using a Carlo Erba NA 1500C/H/N Analyzer coupled to a Thermo Delta V IRMS system (Thermo Fisher Scientific, Waltham, MA, USA). Nitrogen stable isotopes were expressed relative to a standard in δ notation:

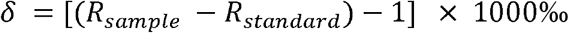

where R represents the ratio of ^15^N to ^14^N. Plants depleted in δ^15^N (i.e., lower δ^15^N) suggests N acquisition primarily from N fixation, while plants enriched in δ^15^N (i.e., higher δ^15^N) suggests N acquisition primarily from the soil (Högberg 1997, Craine et al. 2015). All elemental and stable isotope analyses were conducted at the Stable Isotope Ecology Laboratory at the University of Georgia, USA (http://siel.uga.edu/).

### Landscape metrics

Land use and land cover metrics were calculated for each site to quantify urbanization. We calculated percent impervious surface cover (ISC) manually using Google Earth Pro 7.3.2.5776 (Google Inc., Mountain View, CA, USA) by drawing a 100-m radius around each site and using the polygon tool to draw and measure ISC. We also calculated the normalized difference vegetation index (NDVI), a measure of vegetation cover or site ‘greenness’, for each site from landsat imagery using the “MODISTools” package in R (version 1.1.1; Tuck et al. 2014). We calculated NDVI from 1 June to 31 August at 16-day intervals for each year from 2014-2018 to generate a mean NDVI for each site; NDVI was measured at a spatial resolution of 250 m. Each site received 6 measurements per year for a total of 30 measurements, and we calculated the mean NDVI for each site from these 30 measurements. Finally, for each site, we calculated the distance from the urban center as the distance on an ellipsoid (i.e., geodesic distance) using the “geosphere” package (version 1.5-10; Hijmans 2019); coordinates for the urban center were selected as Toronto City Hall (43.651536°N, -79.383276°W). Distance from the urban center has been shown to be correlated with multiple measures of urbanization in Toronto, and it has been identified as an important predictor of the ecological and evolutionary effects of urbanization for *T. repens* and other systems (Thompson et al. 2016, Johnson et al. 2018, Rivkin et al. 2020).

### Statistical analyses

#### Nodule density mixed-effects models

We used linear mixed-effects models to analyze the response of nodule density against distance from the urban center, percent ISC, and NDVI. We fitted the linear mixed-effects models as:

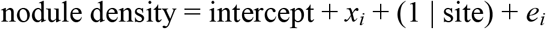

where nodule density was the response, *x*_*i*_ was the fixed effect (distance from the urban center, percent ISC, or NDVI), site was a random effect, and *e*_*i*_ was the residual error associated with the fixed effect. A full model with all predictor variables was not fitted because high multicollinearity between variables (Pearson r > |0.5|) made it difficult to disentangle the effects of individual variables (Supplementary material Appendix 1 Fig. A1). Instead, we used structural equation modelling (see below) to integrate the variables into a single analysis.

All linear mixed-effects models were fitted using the “lme4” (version 1.1-25, Bates et al. 2015) and “lmerTest” (version 3.1-2; Kuznetsova et al. 2017) packages, with models estimated using restricted maximum likelihood. We calculated partial *F*-tests of fixed effects using Type II sums-of-squares, and denominator degrees of freedom were approximated using the Satterthwaite correction for finite sample sizes (Satterthwaite 1946). Response and predictor variables were standardized to a mean of 0 and standard deviation of 1 prior to analyses, and model assumptions were inspected using the DHARMa package (version 0.3.3.0; Hartig 2020). Conditional R^2^, a measure of the variance explained by the fixed and random effects (Nakagawa et al. 2017), was calculated for the models using the “r.squaredGLMM()” function in the “MuMIn” package (version 1.43.17; Barton 2020).

#### White clover δ^15^N and soil N generalized additive models

Changes in white clover δ^15^N and soil N (bulk and adjacent) in response to urbanization were analyzed using generalized additive models (GAMs) using the “mgcv” package (version 1.8-33; Wood 2011, 2017). Generalized additive models are a flexible approach for analyzing data, as non-linear predictors can be fitted with non-parametric smoothing functions to identify effects of the predictors (Hastie and Tibshirani 1986, Wood 2017). We fitted the GAMs as:

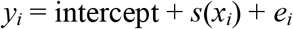

where *y*_*i*_ was the response (white clover δ^15^N, bulk soil N, or adjacent soil N), *x*_*i*_ was the predictor (distance from the urban center, percent ISC, or NDVI), *s* was the non-linear smoothing function for the associated predictor, and *e*_*i*_ was the residual error. Thin-plate regression splines with shrinkage terms (method call: bs = “ts”) were used to smooth all predictors (Wood 2003), and each smooth estimate was constrained to a mean of zero. Residual error was assumed independent with a mean of zero and constant variance. White clover δ^15^N GAMs were fitted to a Gaussian distribution and soil N GAMs were fitted with a logit link to the “betar” distribution, part of the “mgcv” family, to model non-binomial, proportional data. Similar to the nodule density mixed-effects models, high concurvity between variables (Pearson r > |0.5|) precluded the use of a single GAM with all predictor variables (Supplementary material Appendix 1 Fig. A1). Instead, all variables were integrated into the structural equation model (detailed below). All GAMs were estimated using maximum likelihood.

#### Path analysis

A structural equation model (SEM) was constructed to evaluate the direct and indirect causal pathways through which distance from the urban center, percent ISC, NDVI, soil N (bulk and adjacent), nodule density, and white clover δ^15^N interact. Basic univariate and multivariate equations are considered structural if there is sufficient evidence that the predictor or set of predictors has a causal effect on the response (Grace 2006), and SEM is the modelling of a multivariate relationship with two or more structural equations. The robustness of a SEM is based on the overall model fit to the data rather than individual causal pathways within the model (Mitchell 1992, Grace 2006, Grace et al. 2010). Model fit is assessed by comparing expected and observed covariance between predictor and response variables in the SEM using χ^2^ tests (Mitchell 1992, Grace 2006, Grace et al. 2010), and the SEM is considered consistent with the data when expected and observed covariance of the SEM are not different. We fit the hypothesized SEM (Fig. 2), including all causal and correlational pathways, and no removal or addition of causal pathways occurred. Distance from the urban center was fitted as an exogenous variable (i.e., independent variable that affects other variables but is not affected by other variables), while all remaining variables were fitted as endogenous variables (i.e., variables affected by the exogenous variable and that can affect other endogenous variables). Distance from the urban center was fitted to account for extraneous sources of urbanization and environmental variation not explained by the other predictors. All data were standardized by scaling individual variables to a mean of 0 and standard deviation of 1 prior to analysis, and the SEM was estimated using maximum likelihood with robust Satorra-Bentler scaled test statistics (Satorra and Bentler 2001, Rosseel 2012).

Results of the SEM were reported as standardized path coefficients, which show the direction and magnitude of causal pathways between variables and allow for comparison of the strength of relationships within the SEM (Wright 1934, Mitchell 1992, Grace 2006). Direct effects within the SEM are the standardized path coefficient associated with the causal pathway, while indirect effects are quantified by multiplying each path coefficient linking one variable to another within the SEM. For example, in the hypothesized SEM (Fig. 2), an indirect effect of distance from the urban center on bulk soil N is quantified as the path coefficient of distance on percent ISC (percent ISC ∼ distance) times the path coefficient of percent ISC on bulk soil N (bulk soil N ∼ percent ISC). The sign (+ or −) of an indirect effect is the sign of the last causal pathway in the link. Indirect or compounding pathways cannot be calculated across correlational pathways as the direction of the effect is not unidirectional (Wright 1934, Grace 2006).

The SEM was fitted and analyzed using the “lavaan” package (version 0.6-7; Rosseel 2012). All above analyses were conducted using R (version 4.0.2; R Core Team 2020) in the RStudio environment (version 1.4.869; RStudio Team 2020), with data management and figure creation facilitated using the “tidyverse” (version 1.3.0; Wickham et al. 2019).

## Results

### Nodule density, white clover δ^15^N, and soil N

Urbanization had effects on nodule density, white clover δ^15^N, and soil N, although the measure and effect of urbanization varied among response variables. Nodule density increased with increasing distance from the urban center (β = 0.146, SE = 0.066, P = 0.031, R^2^ = 0.147, Table 1, Fig. 3) and NDVI (β = 0.152, SE = 0.064, P = 0.021, R^2^ = 0.146, Table 1, Fig. 3), and nodule density decreased with increasing percent ISC (β = −0.199, SE = 0.063, P = 0.003, R^2^ = 0.151, Table 1, Fig. 3). These effects translated to an increase of (estimate ± SE) 0.006 ± 0.003 in nodules per cm of root (i.e., nodule density) for each 1 km from the city centre, −0.004 ± 0.001 lower nodule density for each 1 percent increase in impervious surface cover, and 0.007 ± 0.003 greater nodule density for each 100 unit increase in NDVI (Supplementary material Appendix 1 Table A1). White clover δ^15^N was strongly and non-linearly predicted by distance from the urban center (*F* = 10.893, P < 0.001, R^2^ = 0.671, deviance explained = 71.0%) but not percent ISC or NDVI (Table 2, Fig. 4). White clover δ^15^N was higher and positive at the urban (0-10 km) and rural (40-47 km) extremes of the gradient, while white clover δ^15^N was lower and negative for the majority of the gradient (10-40 km, Fig. 4). Bulk soil N was predicted by distance from the urban center (χ^2^ = 3.097, P = 0.043, R^2^ = 0.079, deviance explained = 11%; Supplementary material Appendix 1 Table A2, Fig. A2), where bulk soil N decreased with increasing distance from the urban center. Additionally, bulk soil N was weakly predicted by percent ISC (χ^2^ = 1.722, P = 0.098, R^2^ = 0.020, deviance explained = 6%; Supplementary material Appendix 1 Table A2, Fig. A2), where bulk soil N increased with increasing percent ISC; bulk soil N was not predicted by NDVI (Supplementary material Appendix 1 Table A2, Fig. A2); adjacent soil N (i.e., immediately surrounding the plant) was only weakly predicted by percent ISC (χ^2^ = 2.277, P = 0.070, R^2^ = 0.069, deviance explained = 9%; Supplementary material Appendix 1 Table A2, Fig. A2).

**Table 1.** Summary of the nodule density linear mixed-effects model comparing nodule density against distance from the urban center (distance), percent impervious surface cover (ISC), and normalized difference vegetation index (NDVI). We report coefficients (β, slope parameter), standard errors of the coefficients (SE), numerator degrees-of-freedom (NDF), denominator degrees-of-freedom (DDF) approximated following the Satterthwaite method, partial *F*-statistics calculated from Type II sums-of-squares, P-values, and R^2^_conditional_.

| Term | $\beta$ | SE | NDF | DDF | $F$ | P-value | $R^2_{\text{conditional}}$ |
| --- | --- | --- | --- | --- | --- | --- | --- |
| Distance | 0.146 | 0.066 | 1 | 47.723 | 4.949 | 0.031 | 0.147 |
| Percent ISC | -0.199 | 0.063 | 1 | 51.839 | 10.042 | 0.003 | 0.151 |
| NDVI | 0.152 | 0.064 | 1 | 55.268 | 5.640 | 0.021 | 0.146 |

**Figure 3.**
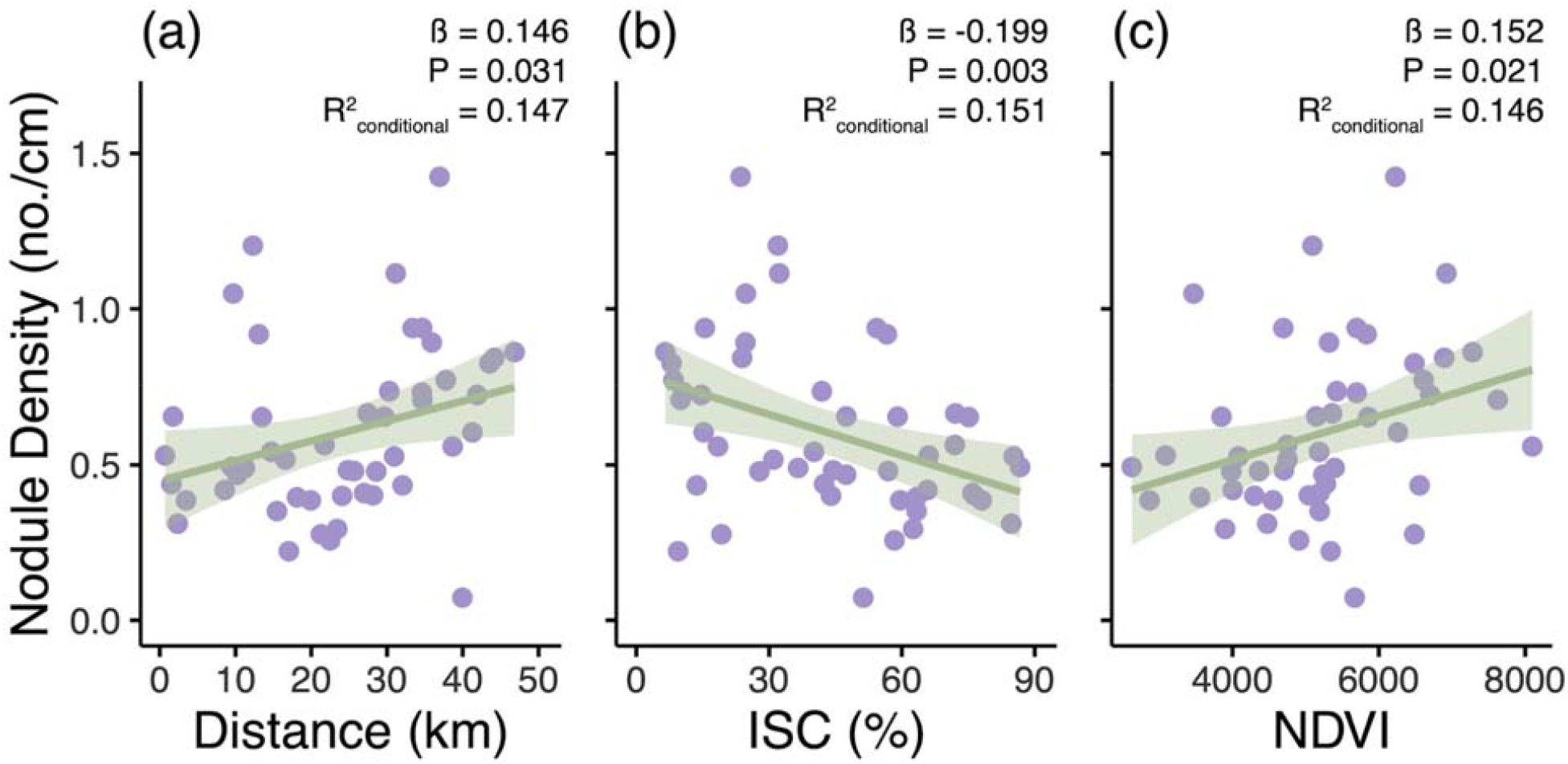
Plots of nodule density against (a) distance from the urban center (Distance), (b) percent impervious surface cover (ISC), and (c) normalized difference vegetation index (NDVI). Lines are lines-of-best-fit (± standard error) from a linear mixed effects model, with the parameter coefficient (β), P-value, and R^2^_conditional_ also provided.

**Table 2.** Summary of the white clover δ^15^N generalized additive models (GAMs) comparing white clover δ^15^N against distance from the urban center (distance), percent impervious surface cover (ISC), and NDVI. We report estimated degrees-of-freedom (EDF), which can differ from 1 because the values are penalized for smoothed parameters; an EDF = 1 would suggest a linear relationship. Reference degrees-of-freedom (REF.df) are used for calculating *F*-statistics and P-values for each smoothed term. We also report the R^2^ and deviance explained for each model, as well as the χ^2^ statistic associated with the smooth term.

| Term | EDF | REF.df | $F$ | P-value | $R^2$ | Deviance Explained |
| --- | --- | --- | --- | --- | --- | --- |
| Distance | 5.654 | 9 | 10.893 | < 0.001 | 0.671 | 71% |
| Percent ISC | < 0.001 | 9 | < 0.001 | 0.820 | $\cong 0.000$ | $\cong 0\%$ |
| NDVI | < 0.001 | 9 | < 0.001 | 0.818 | $\cong 0.000$ | $\cong 0\%$ |

Note: As the white clover $\delta^{15}\text{N}$ GAMs were fitted to a Gaussian error distribution, $F$ -statistics were calculated instead of $\chi^2$ -statistics.

**Figure 4.**
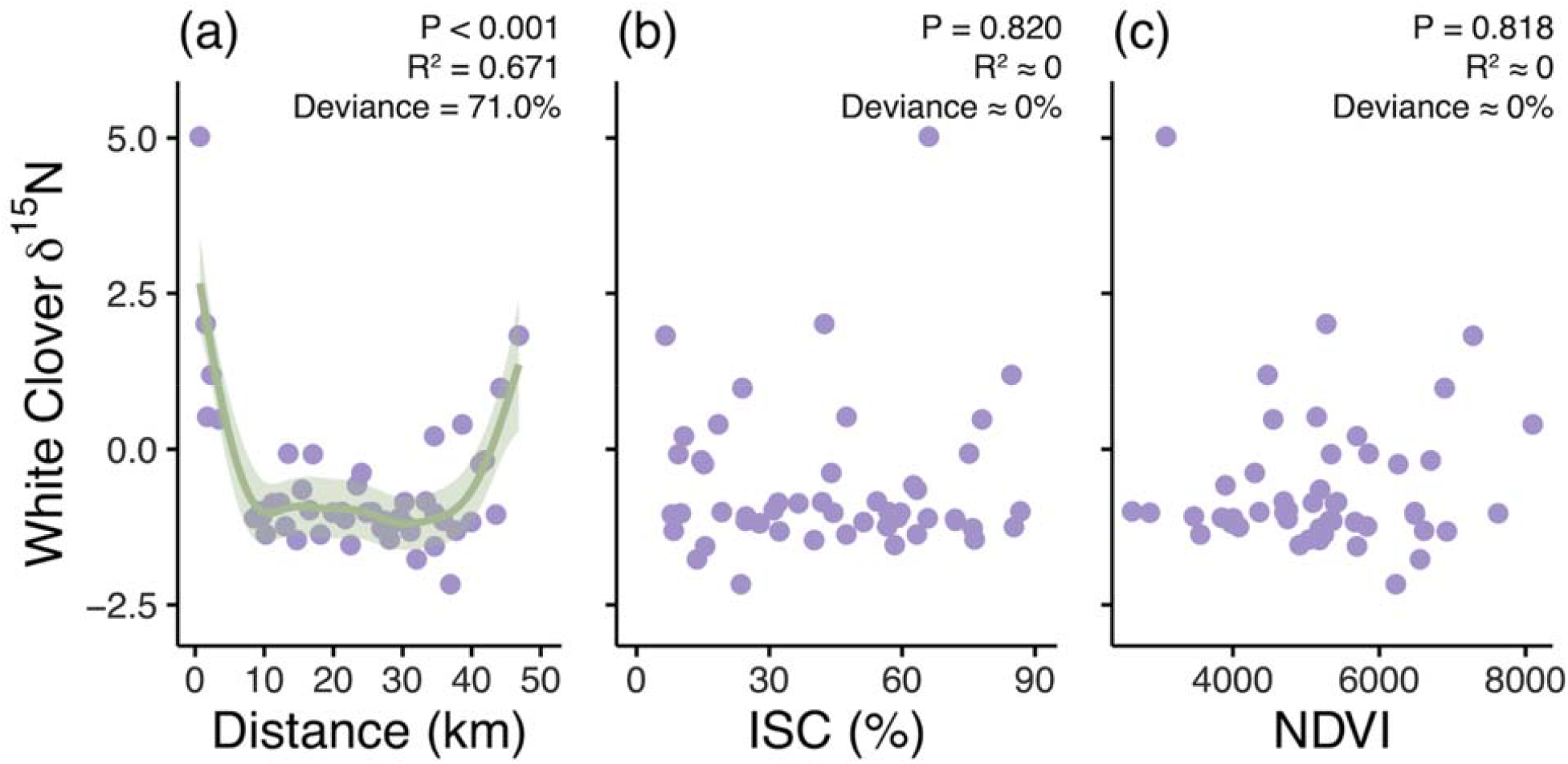
Plots of white clover δ^15^N against (a) distance from the urban center (Distance), (b) percent impervious surface cover (ISC), and (c) normalized difference vegetation index (NDVI). Lines are smoothed curves (± standard error) from a generalized additive model, with the P-value, R^2^, and deviance explained (deviance) also provided. Percent ISC and NDVI were not ecologically-relevant predictors of white clover δ^15^N so lines are not displayed.

### Path analysis

The SEM demonstrated that urbanization had direct and indirect effects on ecosystem structure (i.e., soil N) and the white clover-rhizobium mutualism (Fig. 5). Our hypothesized SEM had good fit between the predicted and observed covariance (χ^2^ = 0.009, df = 1, P = 0.926, Fig. 5). Distance from the urban center had a direct negative relationship with white clover δ^15^N: as the distance from the urban center increased, white clover δ^15^N decreased (path coefficient = −0.60, SE = 0.16, P < 0.001). Bulk soil N decreased with increasing distance from the urban center (path coefficient = −0.29, SE = 0.18, P = 0.099). Further effects of distance from the urban center were mediated through percent ISC on nodule density (percent ISC ∼ distance path coefficient = −0.55, SE = 0.11, P < 0.001; nodule density ∼ percent ISC path coefficient = −0.39, SE = 0.23, P = 0.082; compound path coefficient = |−0.55| x −0.39 = −0.21). Although distance from the urban center was positively related to NDVI (path coefficient = 0.63, SE = 0.08, P < 0.0001), there were no further effects of NDVI in the SEM. Percent ISC and NDVI were negatively correlated (path coefficient = −0.45, SE = 0.13, P = 0.001). Bulk soil N had a negative effect on white clover δ^15^N (path coefficient = −0.23, SE = 0.11, P = 0.040), in which increased bulk soil N decreased white clover δ^15^N. Nodule density and adjacent soil N were negatively correlated (path coefficient = −0.29, SE = 0.11, P = 0.010). Bulk soil N was directly linked to distance from the urban center and then embedded in correlational pathways between adjacent soil N and nodule density (Fig. 5). All path coefficients and associated measures of variation are provided in the supplement (Supplementary material Appendix 1 Table A3).

**Figure 5.**
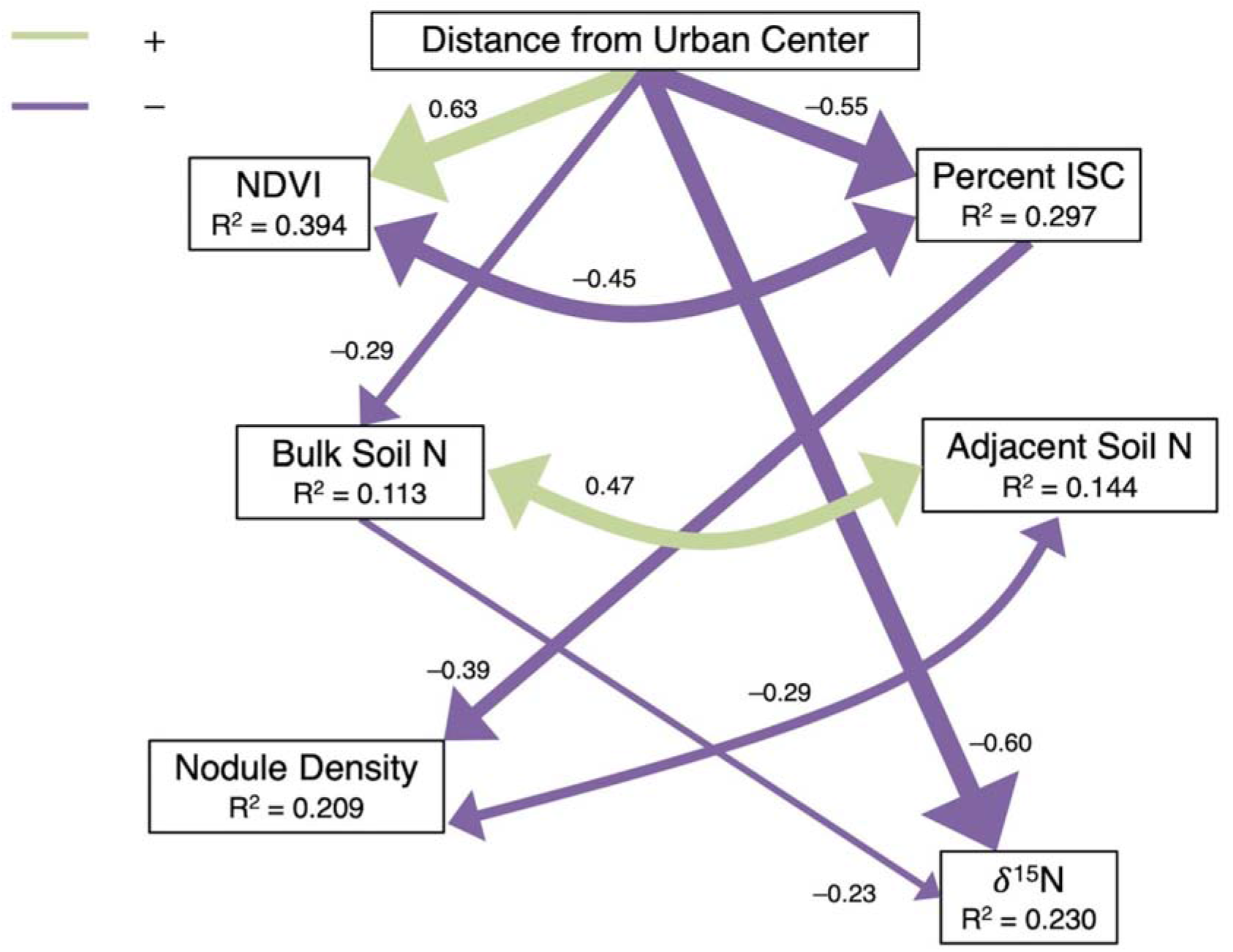
Path diagram showing the causal interactions between distance from the urban center, percent impervious surface cover (ISC), normalized difference vegetation index (NDVI), soil N (bulk and adjacent), nodule density, and white clover δ^15^N. The model had good fit between the predicted and observed covariance (χ^2^ = 0.009, df = 1, P = 0.926). Lines represent influential causal pathways in the model (P < 0.10), with single-headed arrows indicating an unidirectional pathway and double-headed arrows indicating a correlation between variables. Purple lines represent negative pathways, while green lines represent positive pathways; non-influential pathways are not displayed. Standardized path coefficients, which show the direction and magnitude of the relationship between variables, are reported next to each line and line widths are scaled relative to the magnitude of the path coefficients. The R^2^ is reported for each endogenous variable.

## Discussion

Our results show that urbanization alters the ecology of the white clover-rhizobium mutualism and associated patterns in soil nutrients, with support for this conclusion from three key results. First, we observed increased investment in nodules by white clover with decreasing impervious surface cover, increasing NDVI, and greater distance from the urban center (Q1). Second, we found that the source of nitrogen used by white clover varied along the urbanization gradient (Q2). White clover δ^15^N suggested acquisition of N through N-fixing rhizobia for the majority of the urbanization gradient, although increased δ^15^N at the urban and rural extremes suggested direct uptake of N sources from the soil. Finally, we observed direct and indirect effects of urbanization on ecosystem structure and white clover-rhizobium interactions (Q3). Urbanization altered the landscape, changing the amount of green and impervious surface cover surrounding each white clover population, with these changes cascading onto soil N, nodule density, and white clover δ^15^N. Given this evidence, our results suggest that urbanization alters the ecology of the white clover-rhizobia mutualism through direct and indirect pathways.

### Effects of urbanization on the white clover-rhizobium mutualism

Increasing urbanization is frequently associated with increased N deposition and enrichment (Grimm et al. 2008, Zhang et al. 2012), and legumes are expected to have lower investment in rhizobia with greater N availability in the soil (Heath and Tiffin 2007, Heath et al. 2010, Regus et al. 2017). Our results supported this prediction, whereby nodule density, a common measure of investment in mutualistic interactions with rhizobia by host plants (Heath and Tiffin 2007, Heath et al. 2010), increased with decreasing effects of urbanization; these results are supported by other empirical studies in nonurban systems that found reduced investment in nodules with increased N (Heath and Tiffin 2007, Heath et al. 2010, Lau et al. 2012). In our study, nodule density was further influenced by the urban landscape and its effects on the local soil environment. Increasing distance from the urban center was associated with negative effects on bulk soil N and, while nodule density was also negatively related to impervious surface cover, there were entangled dependencies between nodule density, bulk soil N, and adjacent soil N. Given that soil N can regulate nodule formation and development (Streeter and Wong 1988, Omrane and Chiurazzi 2009) and cause shifts in the cost-benefit balance of the white clover-rhizobium mutualism (Lau et al. 2012, Weese et al. 2015, Regus et al. 2017), urban-driven changes to local soil N is a likely driver linking urbanization to altered ecological dynamics in the white clover-rhizobium mutualism.

In addition to nodule density, white clover δ^15^N varied with urbanization. Comparing relative δ^15^N signatures allows for assessing and tracing the source of N used by plants (Högberg 1997, Robinson 2001, Craine et al. 2015). For the majority of the urbanization gradient, white clover δ^15^N was frequently bound between –1‰ and 0‰, suggesting that white clover primarily acquired N through fixation by rhizobia but other sources of N were also used (Högberg 1997, Craine et al. 2015). By contrast, populations at the urban and rural extremes of the gradient exhibited high values of δ^15^N, which suggests decreased reliance on rhizobia for N and increased uptake of N from the soil. A potential explanation for this pattern is the application of fertilizer and management practices. Fertilizer application can directly provide white clover with N, reducing the need for white clover to invest in rhizobia (Vergeer et al. 2008, Heath et al. 2010) and increasing plant δ^15^N (Trammell et al. 2016, 2020). Nitrogen fertilizer use associated with agricultural practices has increased over the past several decades in our study region (Clearwater et al. 2016). Long-term N fertilization can lead to the evolution of less beneficial rhizobia (Weese et al. 2015), which could explain the higher nodule density despite increased white clover δ^15^N (i.e., decreased N fixation) at the agriculturally-intensive rural end of the gradient. This hypothesis will be investigated in future studies.

Although we did not use experimental plant lines to determine the δ^15^N signatures of white clover relying solely on N fixation or on soil acquisition (Högberg 1997, Robinson 2001, Craine et al. 2015), white clover acquiring N solely through fixation by rhizobia has a δ^15^N of – 2‰ to –1‰ (Högberg 1997). Experimentally quantifying the contributions of N fixation and soil uptake along the urbanization gradient would help elucidate the relative importance of these two processes and pools of N. Our existing data show that urbanization is at least partially responsible for variation in nodule density and white clover δ^15^N due to shifts in the costs and benefits of the rhizobia-plant host interaction.

### White clover-rhizobium-soil interactions

Depleted δ^15^N in legumes is associated with increased N fixation by rhizobia, and we expected increased bulk soil N to lead to an increase in white clover δ^15^N (i.e., decreased N fixation). Contrary to our expectation, increased bulk soil N was associated with decreased white clover δ^15^N (i.e., increased N fixation). While contrary to our predictions, increased N can stimulate nodulation, N fixation, and other metabolic processes in rhizobia (Streeter 1985, Simonsen et al. 2015, Forrester and Ashman 2018). We also predicted that lower white clover δ^15^N would increase adjacent soil N as a by-product of N fixation by rhizobia (Hirsch 1992, Poole et al. 2018); this prediction was not supported. While both distance from the urban center and bulk soil N negatively affected white clover δ^15^N, we did not observe a direct link from white clover δ^15^N to adjacent soil N.

We predicted a positive link between nodule density and adjacent soil N as a consequence of increased N fixation, which was expected to enrich soil with N when nodules and plant tissue senesced (Hirsch 1992, Poole et al. 2018). We observed that nodule density was negatively correlated to adjacent soil N. A potential explanation for this response is that white clover produced fewer and larger nodules: larger nodules can convey greater benefits to white clover, increasing N fixation and associated effects on soil N (Porter and Simms 2014, Gano-Cohen et al. 2020). We only measured nodule count and density, so we were unable to test this hypothesis. Our observation of increased adjacent soil N with decreasing nodule density is consistent with inhibition of nodulation due to high N (Streeter and Wong 1988, Omrane and Chiurazzi 2009). Conversely, decreased adjacent soil N with increased nodule densities could also suggest less beneficial rhizobia that fix less N are colonizing roots, which could reduce the enrichment of soil N resulting from the white clover-rhizobium mutualism. Manipulative experiments are needed to disentangle these two possibilities to understand the causal explanation for our observed patterns.

### Urbanization and soil nitrogen

Urbanization had expected effects on soil N, but this was dependent on interactions between white clover and rhizobium and their associated effects on soil and plant N. Bulk soil N was greatest in urban areas, which was consistent with our predictions and observed patterns in other studies (Pouyat et al. 2015, Regus et al. 2017, Trammell et al. 2020). Increased soil N is likely a result of N deposition linked to urbanization (Grimm et al. 2008, Zhang et al. 2012, Regus et al. 2017). Adjacent soil N showed a similar but weaker pattern to bulk soil N in response to impervious surface cover. These results in combination with the SEM show that local soil N is governed by exogenous N inputs (e.g., rain, synthetic fertilizer, pets) into the surrounding soils, although the SEM also suggests a feedback between soil N and rhizobia as bulk and adjacent soil N were correlated, with adjacent soil N further dependent on nodule density. Taken together, these lines of evidence suggest that the effects of urbanization on the white clover-rhizobium mutualism are complex, with local variation in soil N influencing dynamics between white clover, rhizobium, and soil and plant N.

We quantified total soil N to answer our focal questions for this study, but other measures of soil N can be important to consider. For example, total N does not discriminate between different types of N (e.g., NO_3_, NO_2_, and NH_3_). Identifying the type and quantifying the relative amounts of each species of N could provide a clearer link between urbanization and N deposition, as different forms of N are important for both plant and rhizobia physiology and metabolism (Wallsgrove et al. 1983). In addition to further investigation of soil N content, soil δ^15^N could integrate inputs, metabolic processes, and transformations of N in ecosystems (Robinson 2001). For example, a typical fertilizer has a δ^15^N around 0‰ and other anthropogenic sources have enriched δ^15^N signatures (Robinson 2001, Michener and Lajtha 2007), which can be reflected in the tissues of organisms utilizing these sources (Robinson 2001, Trammell et al. 2016). Urban environments frequently have increased and less variable soil δ^15^N relative to nearby nonurban or rural environments (Trammell et al. 2020). Therefore, using soil δ^15^N could also help to identify a direct link from urbanization to changes in soil N and ultimately legume-rhizobia mutualisms.

### Limitations

Our study had several limitations that contextualize our conclusions. First, we focused on the focal white clover-rhizobia mutualism without considering how other coexisting plants could affect the focal mutualism and soil nutrient patterns. White clover was frequently collected in patches of grass, especially *Poa annua*, and near other legumes, predominantly *Medicago lupulina* and occasionally other *Trifolium* species (*T. pratense, T. hybridum*; DMS, personal observation). White clover competes for space and nutrients with these other plants and this competition could have affected the response of the focal mutualism to urbanization, especially if plant species composition changed along the urbanization gradient as has been reported elsewhere (Hope et al. 2003, Knapp et al. 2008, 2012). Other legumes might have also affected the observed patterns in soil N, although we did explicitly avoid collecting soil from patches of other legume species so this possibility is unlikely. Second, we did not investigate how the microbial community beyond rhizobium in the soil and in the roots of white clover varies with urbanization. Microbes are crucial for plant community assembly and responses to biotic and abiotic stress (van der Heijden et al. 2008, Fitzpatrick et al. 2018). Additionally, the ostensibly pairwise mutualistic interaction between white clover and rhizobia can be altered in the presence of other bacteria or fungi (Heath and Tiffin 2007, García Parisi et al. 2015). With recent studies documenting changes in microbial diversity and composition in response to urbanization (Xu et al. 2014, Reese et al. 2016), it is plausible that variation in microbial communities in the mosaic of urban environments could alter the ecological effects of the white clover-rhizobium interaction and shift the cost-benefit balance of the mutualism (García Parisi et al. 2015, Burghardt et al. 2018, Batstone et al. 2020). Lastly, temporal variation in N deposition could have a differential effect on soil N along the urbanization gradient (Zbieranowski and Aherne 2011, 2012), with rural soil N being more variable than urban soil N (Trammell et al. 2020). As we only measured white clover δ^15^N, we were only able to identify the N source integrated into plant tissue over the growing season and account for temporal variation in N sources. Measuring soil δ^15^N would have traced potential changes to N inputs in the system, but temporal variation in soil N and linkages to a legume-rhizobia mutualism was outside the scope of this study. Notwithstanding these limitations, our study shows that urbanization alters the ecological and ecosystem-level effects of the white clover-rhizobia mutualism.

### Conclusion

Our study represents an evaluation of the effects of urbanization on an ecologically-important mutualism. To date, urban ecological research has principally evaluated species interactions by focusing on plant-pollinator and plant-herbivore interactions (Youngsteadt et al. 2015, Aronson et al. 2016, Harrison et al. 2018, Miles et al. 2019, Rivkin et al. 2020). We have extended this field by investigating the mutualistic interaction between a plant (white clover) and its mutualistic, microbial symbionts (rhizobia). White clover invested in more nodules and relied on rhizobia for N fixation with less urbanization, and increased urbanization directly and negatively affected investment in rhizobia by white clover, with soil N playing a critical role linking urbanization to the mutualistic interaction between white clover and rhizobia. In conclusion, our results demonstrate the direct and indirect effects of urbanization on the cost-benefit balance and ecological consequences of a legume-rhizobium mutualism.

## Data Accessibility Statement

Data and code are available on Zenodo <http://doi.org/10.5281/zenodo.4459724> (Murray-Stoker and Johnson 2021).

## Acknowledgements

We thank S. Munim and K. Murray-Stoker for assistance with field work and C. Sastropranoto for help with soil sample processing. We thank R. Rivkin, J. Santangelo, L. Albano, S. Breitbart, S. Koch, and L. Miles for providing comments on earlier drafts of the manuscript.

## Author Contributions

<u>David Murray-Stoker:</u> Conceptualization (Equal); Data curation (Lead); Formal analysis (Lead); Investigation (Equal); Methodology (Equal); Writing-original draft (Lead); Writing-review & editing (Equal)

<u>Marc T. J. Johnson:</u> Conceptualization (Equal); Data curation (Supporting); Formal analysis (Supporting); Funding acquisition (Lead); Investigation (Equal); Methodology (Equal); Writing-original draft (Supporting); Writing-review & editing (Equal)

## References

Andrews, M. and Andrews, M. E. 2017. Specificity in legume-rhizobia symbioses. - Int. J. Mol. Sci. 18: 705.

Aronson, M. F. J. et al. 2016. Hierarchical filters determine community assembly of urban species pools. - Ecology 97: 2952–2963.

Baker, M. J. and Williams, W. M. 1987. White Clover. - C.A.B. International.

Barrett, J. P. and Silander, J. A. 1992. Seedling recruitment limitation in white clover (Trifolium repens; Leguminosae). - Am. J. Bot. 79: 643–649.

Barton, K. 2020. MuMIn: Multi-Model Inference. - R package version 1.43.17. <https://CRAN.R-project.org/package=MuMIn>

Bates, D. et al. 2015. Fitting linear mixed-effects models using lme4. - J. Stat. Softw. 67: 1–48.

Batstone, R. T. et al. 2020. Experimental evolution makes microbes more cooperative with their local host genotype. - Science 370: 476–478.

Burghardt, L. T. et al. 2018. Select and resequence reveals relative fitness of bacteria in symbiotic and free-living environments. - Proc. Natl. Acad. Sci. U. S. A. 115: 2425– 2430.

Clearwater, R. L. et al. 2016. Environmental sustainability of Canadian agriculture: Agri-environmental indicator report series-Report #4. - Agriculture and Agri-Food Canada.

Cleland, E. E. and Harpole, W. S. 2010. Nitrogen enrichment and plant communities. - Ann. N. Y. Acad. Sci. 1195: 46–61.

Craine, J. M. et al. 2015. Ecological interpretations of nitrogen isotope ratios of terrestrial plants and soils. - Plant Soil 396: 1–26.

De León, L. F. et al. 2019. Urbanization erodes niche segregation in Darwin’s finches. - Evol. Appl. 12: 1329–1343.

Fitzpatrick, C. R. et al. 2018. Assembly and ecological function of the root microbiome across angiosperm plant species. - Proc. Natl. Acad. Sci. U. S. A. 115: E1157–E1165.

Forrester, N. J. and Ashman, T.-L. 2018. Nitrogen fertilization differentially enhances nodulation and host growth of two invasive legume species in an urban environment. - J. Urban Ecol. 4: juy021.

Galloway, J. N. et al. 2003. The nitrogen cascade. - BioScience 53: 341.

Gano-Cohen, K. A. et al. 2020. Recurrent mutualism breakdown events in a legume rhizobia metapopulation. - Proc. R. Soc. B Biol. Sci. 287: 20192549.

García Parisi, P.A. et al. 2015. Multi-symbiotic systems: functional implications of the coexistence of grass-endophyte and legume-rhizobia symbioses. - Oikos 124: 553–560.

Grace, J. 2006. Structural Equation Modeling and Natural Systems. - Cambridge University Press.

Grace, J. B. et al. 2010. On the specification of structural equation models for ecological systems. - Ecol. Monogr. 80: 67–87.

Grimm, N. B. et al. 2008. Global change and the ecology of cities. - Science 319: 756–760.

Groffman, P. M. et al. 2014. Ecological homogenization of urban USA. - Front. Ecol. Environ. 12: 74–81.

Harrison, J. G. et al. 2018. Deconstruction of a plant-arthropod community reveals influential plant traits with nonlinear effects on arthropod assemblages. - Funct. Ecol. 32: 1317– 1328.

Hartig, F. 2020. DHARMa: Residual Diagnostics for Hierarchical (Multi-Level / Mixed) Regression Models. - R package version 0.3.3.0. <https://CRAN.R-project.org/package=DHARMa>

Hastie, T. and Tibshirani, R. 1986. Generalized additive models. - Stat. Sci. 1: 291–310.

Heath, K. D. and Tiffin, P. 2007. Context dependence in the coevolution of plant and rhizobial mutualists. - Proc. R. Soc. B Biol. Sci. 274: 1905–1912.

Heath, K. D. et al. 2010. Mutualism variation in the nodulation response to nitrate. - J. Evol. Biol. 23: 2494–2500.

Hijmans, R. J. 2019. geosphere: Spherical Trigonometry. - R package version 1.5-10. <https://CRAN.R-project.org/package=geosphere>

Hirsch, A. M. 1992. Developmental biology of legume nodulation. - New Phytol. 122: 211–237.

Högberg, P. 1997. 15N natural abundance in soil-plant systems. - New Phytol. 137: 179–203.

Hope, D. et al. 2003. Socioeconomics drive urban plant diversity. - Proc. Natl. Acad. Sci. U. S. A. 100: 8788–8792.

Irwin, R. E. et al. 2014. Plant–animal interactions in suburban environments: implications for floral evolution. - Oecologia 174: 803–815.

Johnson, M. T. J. et al. 2018. Contrasting the effects of natural selection, genetic drift and gene flow on urban evolution in white clover (Trifolium repens). - Proc. R. Soc. B Biol. Sci. 285: 20181019.

Kaye, J. et al. 2006. A distinct urban biogeochemistry? - Trends Ecol. Evol. 21: 192–199.

Kemball, W. D. and Marshall, C. 1995. Clonal integration between parent and branch stolons in white clover: a developmental study. - New Phytol. 129: 513–521.

Knapp, S. et al. 2008. Challenging urban species diversity: contrasting phylogenetic patterns across plant functional groups in Germany. - Ecol. Lett. 11: 1054–1064.

Knapp, S. et al. 2012. Phylogenetic and functional characteristics of household yard floras and their changes along an urbanization gradient. - Ecology 93: S83–S98.

Kuznetsova, A. et al. 2017. lmerTest package: tests in linear mixed effects models. - J. Stat. Softw. 82: 1–26.

Lau, J. A. et al. 2012. Direct and interactive effects of light and nutrients on the legume-rhizobia mutualism. - Acta Oecologica 39: 80–86.

Leibold, M. A. and Chase, J. M. 2017. Metacommunity Ecology. - Princeton University Press.

Martínez-Romero, E. and Caballero-Mellado, J. 1996. Rhizobium phylogenies and bacterial genetic diversity. - Crit. Rev. Plant Sci. 15: 113–140.

McKinney, M. L. 2002. Urbanization, biodiversity, and conservation. - BioScience 52: 883–890.

Michener, R. and Lajtha, K. 2007. Stable Isotopes in Ecology and Environmental Science. -Blackwell Publishing.

Miles, L. S. et al. 2019. Urbanization shapes the ecology and evolution of plant-arthropod herbivore interactions. - Front. Ecol. Evol. 7: 310.

Mitchell, R. J. 1992. Testing evolutionary and ecological hypotheses using path analysis and structural equation modelling. - Funct. Ecol. 6: 123–129.

Murray-Stoker, D., and M Johnson. 2021. Data from: dmurraystoker/TRhizo-urbanTerreN: Ecological consequences of urbanization on a legume-rhizobia mutualism (Version V1.0). - Zenodo <http://doi.org/10.5281/zenodo.4459724>

Nakagawa, S. et al. 2017. The coefficient of determination R^2^ and intra-class correlation coefficient from generalized linear mixed-effects models revisited and expanded. - J. R. Soc. Interface 14: 20170213.

Omrane, S. and Chiurazzi, M. 2009. A variety of regulatory mechanisms are involved in the nitrogen-dependent modulation of the nodule organogenesis program in legume roots. -Plant Signal. Behav. 4: 1066–1068.

Parsons, A. W. et al. 2019. Urbanization focuses carnivore activity in remaining natural habitats, increasing species interactions. - J. Appl. Ecol. 56: 1894–1904.

Poole, P. et al. 2018. Rhizobia: from saprophytes to endosymbionts. - Nat. Rev. Microbiol. 16: 291–303.

Porter, S. S. and Simms, E. L. 2014. Selection for cheating across disparate environments in the legume-rhizobium mutualism. - Ecol. Lett. 17: 1121–1129.

Pouyat, R. V. et al. 2015. A global comparison of surface soil characteristics across five cities: a test of the urban ecosystem convergence hypothesis. - Soil Sci. 180: 136–145.

R Core Team 2020. R: a language and environment for statistical computing. - version 4.0.3. <https://cran.r-project.org/>

R Studio Team 2020. RStudio: Integrated development for R. - version 1.4.869. <https://www.rstudio.com/>

Raupp, M. J. et al. 2010. Ecology of herbivorous arthropods in urban landscapes. - Annu. Rev. Entomol. 55: 19–38.

Reese, A. T. et al. 2016. Urban stress is associated with variation in microbial species composition—but not richness—in Manhattan. - ISME J. 10: 751–760.

Regus, J. U. et al. 2017. Nitrogen deposition decreases the benefits of symbiosis in a native legume. - Plant Soil 414: 159–170.

Rivkin, L. R. et al. 2020. Variation in pollinator-mediated plant reproduction across an urbanization gradient. - Oecologia 192: 1073–1083.

Robinson, D. 2001. δ^15^N as an integrator of the nitrogen cycle. - Trends Ecol. Evol. 16: 153–162.

Rocha, E. A. and Fellowes, M. D. E. 2018. Does urbanization explain differences in interactions between an insect herbivore and its natural enemies and mutualists? - Urban Ecosyst. 21: 405–417.

Rosseel, Y. 2012. lavaan: an R package for structural equation modeling. - J. Stat. Softw. 48: 1– 36.

Santangelo, J. S. et al. 2020. Multivariate phenotypic divergence along an urbanization gradient. - Biol. Lett. 16: 20200511.

Satorra, A. and Bentler, P. M. 2001. A scaled difference chi-square test statistic for moment structure analysis. - Psychometrika 66: 507–514.

Satterthwaite, F. E. 1946. An approximate distribution of estimates of variance components. - Biom. Bull. 2: 110–114.

Seto, K. C. et al. 2010. The new geography of contemporary urbanization and the environment. - Annu. Rev. Environ. Resour. 35: 167–194.

Simonsen, A. K. et al. 2015. Short-term fertilizer application alters phenotypic traits of symbiotic nitrogen fixing bacteria. - PeerJ 3: e1291.

Stevens, C. J. et al. 2018. Atmospheric nitrogen deposition in terrestrial ecosystems: Its impact on plant communities and consequences across trophic levels. - Funct. Ecol. 32: 1757– 1769.

Streeter, J. G. 1985. Nitrate inhibition of legume nodule growth and activity. - Plant Physiol. 77: 325–328.

Streeter, J. and Wong, P. P. 1988. Inhibition of legume nodule formation and N_2_ fixation by nitrate. - Crit. Rev. Plant Sci. 7: 1–23.

Thompson, K. A. et al. 2016. Urbanization drives the evolution of parallel clines in plant populations. - Proc. R. Soc. B Biol. Sci. 283: 20162180.

Thomson, D. M. and Page, M. L. 2020. The importance of competition between insect pollinators in the Anthropocene. - Curr. Opin. Insect Sci. 38: 55–62.

Trammell, T. L. E. et al. 2016. Plant nitrogen concentration and isotopic composition in residential lawns across seven US cities. - Oecologia 181: 271–285.

Trammell, T. L. E. et al. 2020. Urban soil carbon and nitrogen converge at a continental scale. - Ecol. Monogr. 90: e01401.

Tuck, S. L. et al. 2014. MODISTools - downloading and processing MODIS remotely sensed data in R. - Ecol. Evol. 4: 4658–4668.

van der Heijden, M. G. A. et al. 2008. The unseen majority: soil microbes as drivers of plant diversity and productivity in terrestrial ecosystems. - Ecol. Lett. 11: 296–310.

Vergeer, P. et al. 2008. Geographical variation in the response to nitrogen deposition in Arabidopsis lyrata petraea. - New Phytol. 179: 129–141.

Wallsgrove, R. M. et al. 1983. Photosynthesis, photorespiration and nitrogen metabolism. - Plant Cell Environ. 6: 301–309.

Weese, D. J. et al. 2015. Long-term nitrogen addition causes the evolution of less-cooperative mutualists. - Evolution 69: 631–642.

Wickham, H. et al. 2019. Welcome to the tidyverse. - J. Open Source Softw. 4: 1686.

Williams, N. S. G. et al. 2009. A conceptual framework for predicting the effects of urban environments on floras. - J. Ecol. 97: 4–9.

Wisz, M. S. et al. 2013. The role of biotic interactions in shaping distributions and realised assemblages of species: implications for species distribution modelling. - Biol. Rev. 88: 15–30.

Wood, S. N. 2003. Thin plate regression splines. - J. R. Stat. Soc. Ser. B Stat. Methodol. 65: 95– 114.

Wood, S. N. 2011. Fast stable restricted maximum likelihood and marginal likelihood estimation of semiparametric generalized linear models. - J. R. Stat. Soc. Ser. B Stat. Methodol. 73: 3–36.

Wood, S. N. 2017. Generalized Additive Models: An Introduction with R, Second Edition. - Chapman and Hall/CRC.

Wright, S. 1934. The method of path coefficients. - Ann. Math. Stat. 5: 161–215.

Xu, H.-J. et al. 2014. Does urbanization shape bacterial community composition in urban park soils? A case study in 16 representative Chinese cities based on the pyrosequencing method. - FEMS Microbiol. Ecol. 87: 182–192.

Youngsteadt, E. et al. 2015. Do cities simulate climate change? A comparison of herbivore response to urban and global warming. - Glob. Change Biol. 21: 97–105.

Zbieranowski, A. L. and Aherne, J. 2011. Long-term trends in atmospheric reactive nitrogen across Canada: 1988–2007. - Atmos. Environ. 45: 5853–5862.

Zbieranowski, A. L. and Aherne, J. 2012. Ambient concentrations of atmospheric ammonia, nitrogen dioxide and nitric acid across a rural–urban–agricultural transect in southern Ontario, Canada. - Atmos. Environ. 62: 481–491.

Zhang, L. et al. 2012. Nitrogen deposition to the United States: distribution, sources, and processes. - Atmospheric Chem. Phys. 12: 4539–4554.

Zheng, M. et al. 2019. Global pattern and controls of biological nitrogen fixation under nutrient enrichment: A metaLJanalysis. - Glob. Change Biol. 25: 3018–3030.

Ziter, C. 2016. The biodiversity-ecosystem service relationship in urban areas: a quantitative review. - Oikos 125: 761–768.

