## Appendix 1 for "Ecological consequences of urbanization on a legume-rhizobia mutualism"

**Table A1.** Summary of the unscaled nodule density mixed-effects model comparing nodule density against distance from the urban center (distance), percent impervious surface cover (ISC), and normalized difference vegetation index (NDVI). We report model coefficients (β, slope parameter), standard errors of the coefficients (SE), numerator degrees-of-freedom (NDF), denominator degrees of freedom (DDF) approximated following the Satterthwaite method, *F*-statistics calculated from Type II sums-of-squares, P-values, and $R^2_{conditional}$.

| Term | β | SE | NDF | DDF | F | P-value | $R^2_{conditional}$ |
|---|---|---|---|---|---|---|---|
| Distance | 0.006 | 0.003 | 1 | 46.614 | 4.612 | 0.037 | 0.167 |
| Percent ISC | −0.004 | 0.001 | 1 | 46.688 | 8.688 | 0.005 | 0.166 |
| NDVI | < 0.001 | < 0.001 | 1 | 46.518 | 5.185 | 0.027 | 0.167 |

**Table A2.** Summary of the plant bulk and adjacent soil N generalized additive models (GAMs) comparing plant soil N against distance from the urban center (distance), percent impervious surface cover (ISC), and normalized difference vegetation index (NDVI). We report estimated degrees of freedom (EDF), which can differ from 1 because the values are penalized for smoothed parameters; an EDF = 1 would suggest a linear relationship. Reference degrees of freedom (REF.df) are used for calculating $\chi^2$-statistics and P-values for each smoothed term. We also report the $R^2$ and deviance explained for each model.

| Term | EDF | REF.df | $\chi^2$ | P-value | $R^2$ | Deviance Explained |
|---|---|---|---|---|---|---|
| **Bulk Soil N** | | | | | | |
| Distance | 0.765 | 9 | 3.097 | 0.043 | 0.079 | 11% |
| Percent ISC | 0.635 | 9 | 1.722 | 0.098 | 0.020 | 6% |
| NDVI | 0.247 | 9 | 0.319 | 0.255 | 0.011 | 2% |
| **Adjacent Soil N** | | | | | | |
| Distance | 0.121 | 9 | 0.144 | 0.275 | 0.003 | 1% |
| Percent ISC | 0.698 | 9 | 2.277 | 0.070 | 0.069 | 9% |
| NDVI | 0.475 | 9 | 0.872 | 0.175 | 0.022 | 4% |

Note: $\chi^2$-statistics were calculated instead of *F*-statistics as the soil N GAMs were fitted to a beta distribution (“betar” in the “mgcv” package in R) for non-binomial, proportional data.

**Table A3.** Path coefficients for each causal and correlational pathway in the SEM. We report the structural equation (causal pathways indicated with ~ and correlational pathways indicated with ~~), path coefficient, standard error of the path coefficient (SE), *z*-statistic, P-value, and 95% confidence interval (95% CI) of the path coefficient.

| Structural Equation | Path Coefficient | SE | *z* | P-value | 95% CI |
|---|---|---|---|---|---|
| ISC ~ Distance | −0.55 | 0.11 | −5.207 | < 0.001 | −0.75, −0.34 |
| NDVI ~ Distance | 0.63 | 0.08 | 7.432 | < 0.001 | 0.46, 0.79 |
| Bulk N ~ Distance | −0.29 | 0.18 | −1.649 | 0.099 | −0.63, −0.05 |
| Bulk N ~ ISC | 0.07 | 0.28 | 0.241 | 0.809 | −0.47, 0.60 |
| Bulk N ~ NDVI | −0.01 | 0.26 | −0.034 | 0.973 | −0.52, 50 |
| Adjacent N ~ ISC | 0.32 | 0.23 | 1.375 | 0.169 | −0.14, 0.78 |
| Adjacent N ~ NDVI | −0.03 | 0.22 | −0.123 | 0.902 | −0.46, 0.41 |
| Adjacent N ~ Clover $\delta^{15}N$ | −0.14 | 0.10 | −1.396 | 0.163 | −0.33, 0.05 |
| Nod. Density ~ Distance | 0.14 | 0.20 | 0.695 | 0.487 | −0.25. 0.53 |
| Nod. Density ~ ISC | −0.39 | 0.23 | -−1.742 | 0.082 | −0.83, 0.05 |
| Nod. Density ~ NDVI | −0.04 | 0.17 | −0.242 | 0.809 | −0.37, 0.29 |
| Nod. Density ~ Bulk N | −0.01 | 0.17 | −0.040 | 0.968 | −0.33, 0.32 |
| Clover $\delta^{15}N$ ~ Distance | −0.60 | 0.16 | −3.840 | < 0.001 | −0.90, −0.29 |
| Clover $\delta^{15}N$ ~ ISC | −0.08 | 0.11 | −0.731 | 0.464 | −0.31, 0.14 |
| Clover $\delta^{15}N$ ~ NDVI | 0.24 | 0.21 | 1.140 | 0.254 | −0.17, 0.64 |
| Clover $\delta^{15}N$ ~ Bulk N | −0.23 | 0.11 | −2.059 | 0.040 | −0.44, −0.01 |
| Clover $\delta^{15}N$ ~ Nod. Density | −0.10 | 0.14 | −0.708 | 0.479 | −0.37, 0.17 |
| Bulk N ~~ Adjacent N | 0.47 | 0.14 | 3.447 | < 0.001 | 0.20, 0.73 |
| ISC ~~ NDVI | −0.45 | 0.12 | −3.797 | < 0.001 | −0.68, −0.22 |
| Adjacent N ~~ Nod. Density | −0.29 | 0.11 | −2.588 | 0.010 | −0.52, −0.07 |

Note: Variables are abbreviated as: distance from the urban center = Distance; percent impervious surface cover = ISC; normalized difference vegetation index = NDVI; bulk soil N = Bulk N; adjacent soil N = Adjacent N; nodule density = Nod. Density; and white clover $\delta^{15}N$ = Clover $\delta^{15}N$.

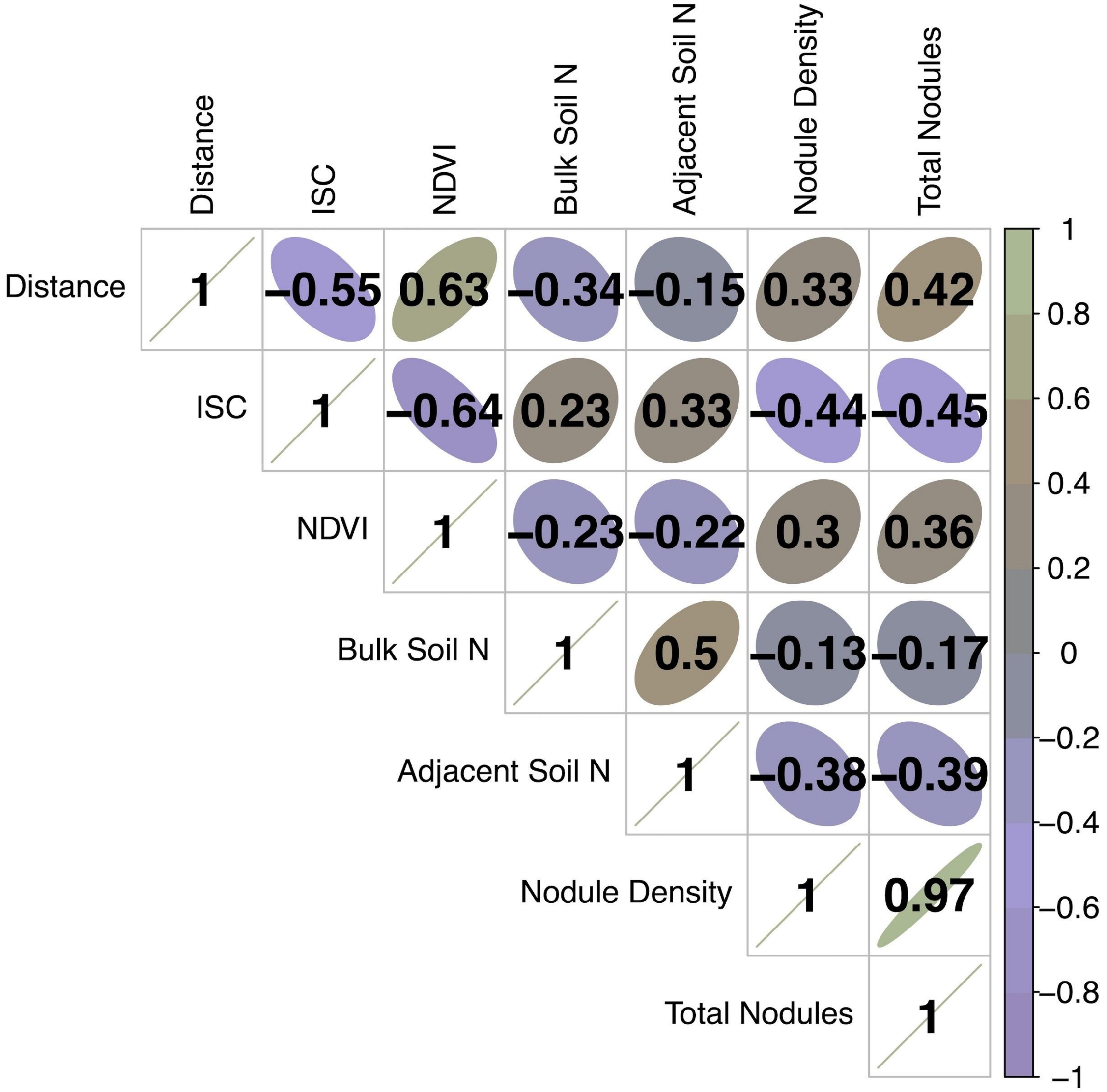


**Figure A1.** Correlations between all variables in the dataset. We calculated all pairwise Pearson correlation coefficients between distance from the urban center (distance), percent impervious surface cover (ISC), normalized difference vegetation index (NDVI), bulk soil N, adjacent soil N, nodule density, and nodule count (total nodules).

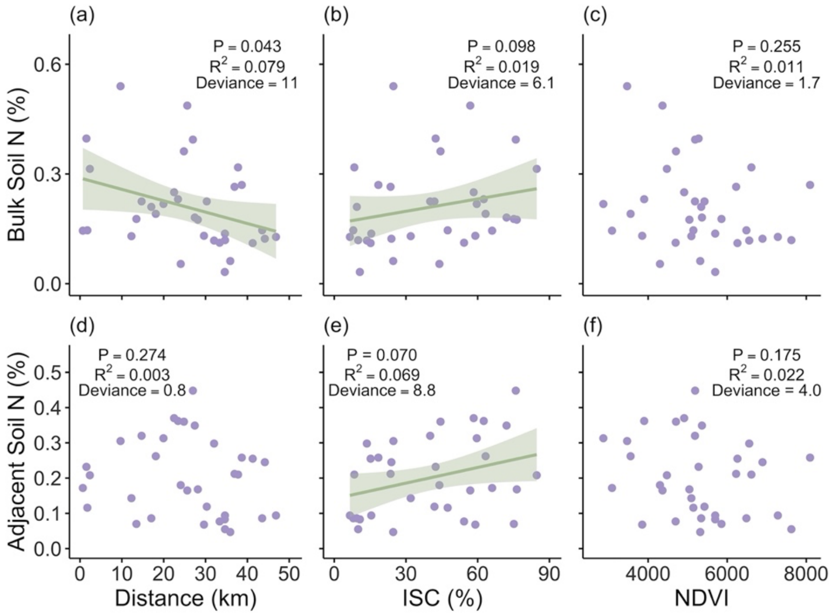


**Figure A2.** Plots of bulk soil N (a, b, c) and adjacent soil N (d, e, f) against distance from the urban center (Distance; a, d), percent impervious surface cover (ISC; b, e), and normalized difference vegetation index (NDVI; c, f). Lines are smoothed curves (± standard error) from a generalized additive model, with the P-value, $R^2$, and deviance explained (deviance) also provided. Distance was not an ecologically-relevant predictor for adjacent soil N and NDVI was not a relevant predictor for either bulk soil N or adjacent soil N, so lines are not displayed.
